# Identification of tail-binding proteins of Arabidopsis class VIII myosin ATM1 using TurboID proximity labeling and AlphaFold3

**DOI:** 10.64898/2026.04.22.720059

**Authors:** Sakura Nagata, Shun Sakuraba, Emi Mishiro-Sato, Takashi L. Shimada, Yuzuki Oe, Kohei Tachibana, Jun Obara, Motoki Tominaga, Kohji Ito, Takeshi Haraguchi

**Author notes:** Corresponding authors: Takeshi Haraguchi, E-mail. **Footnotes:** The mass spectrometry proteomics data reported in this paper have been deposited to the jPOST repository (Okuda et al. 2025) with the dataset identifiers JPST004539 and PXD076613.

## Abstract

Higher plants possess two classes of myosin molecular motors, class XI and class VIII, both unique to the plant lineage. The diverse cellular functions of class XI myosins, including organelle transport and nuclear positioning, have been elucidated largely through **s**ystematic identification of cargo adaptor proteins that bind to their globular tail domains (GTDs). In contrast, no proteome-wide screen for class VIII myosin tail-binding proteins has been reported; the few known interacting proteins were each discovered through studies focused on the binding partner rather than on the myosin itself, leaving the full repertoire of class VIII myosin-associated proteins largely unknown. Here, we employed TurboID-based proximity labeling to systematically identify proteins associated with the GTD of the class VIII myosin ATM1 in *Arabidopsis thaliana*, as this approach covalently biotinylates neighboring proteins *in vivo*, enabling their identification even after proteolytic degradation during cell lysis. We identified 233 non-redundant candidate ATM1-proximal proteins. Candidates were prioritized by AlphaFold3-based protein complex structure prediction and validated by co-immunoprecipitation. We identified two ATM1-associated proteins: C3H61/AtTZF5, a tandem zinc finger protein involved in mRNA turnover at processing bodies and stress granules; and SFH7, a Sec14-nodulin domain protein that mediates phosphatidic acid transfer from the endoplasmic reticulum to chloroplasts. These findings provide initial evidence linking ATM1 to proteins involved in post-transcriptional gene regulation and interorganellar lipid transport, raising the possibility of previously unrecognized connections between class VIII myosins and these cellular processes.

## Introduction

Myosin is a motor protein that converts chemical energy liberated by ATP hydrolysis into directed movement on actin filaments. Phylogenetic analyses of myosin motor domain sequences have identified at least 79 myosin classes throughout eukaryotes (Kollmar and Muhlhausen 2017). Plants possess only two classes of myosins, class VIII and class XI, both of which are found exclusively in plants (Reddy and Day 2001; Tominaga and Nakano 2012). Class XI myosins are related to fungal and animal class V myosins, whereas class VIII myosins represent a distinct evolutionary lineage with no clear counterparts in other kingdoms (Odronitz and Kollmar 2007).

Class XI myosins have been extensively characterized as the primary driving force for cytoplasmic streaming. Arabidopsis contains 13 class XI myosin genes with diverse expression patterns and enzymatic properties (Haraguchi et al. 2018; Peremyslov et al. 2011). Their in vitro sliding velocities vary widely, with estimated values ranging from >10 μm/s for pollen-specific myosins to ∼0.5 μm/s for the nuclear envelope-associated myosin XI-I (Haraguchi et al. 2018). Beyond these differences in expression and motility, the functional roles of individual class XI myosins have been elucidated largely through the identification of tail-binding proteins that serve as cargo adaptors. MyoB family proteins recruit class XI myosins to endomembrane vesicles (Peremyslov et al. 2013), MadA and MadB mediate root hair polar growth (Kurth et al. 2017), and WIT1/WIT2 anchor myosin XI-I to the nuclear envelope (Tamura et al. 2013). Class XI myosin mutants exhibit pronounced defects in organelle transport and cell expansion (Peremyslov et al. 2010). Thus, identification of tail-binding proteins has been instrumental in understanding the diverse functions of class XI myosins.

In contrast, the physiological functions of class VIII myosins remain poorly understood, despite considerable advances in characterizing their enzymatic properties and subcellular localization. Arabidopsis contains four class VIII myosin genes (ATM1, ATM2, VIIIA, and VIIIB) (Reddy and Day 2001). ATM1 localizes to plasmodesmata, the cortical endoplasmic reticulum, and endosomes (Golomb et al. 2008; Haraguchi et al. 2014), and exhibits enzymatic properties markedly different from class XI myosins: slow ATPase activity (V_max_ ∼4 s⁻¹), low sliding velocity (∼0.2 μm/s), and an extremely high duty ratio (∼90%), suggesting that ATM1 functions as a molecular anchor or tether rather than a processive transporter (Haraguchi et al. 2014). However, the quadruple mutant lacking all four class VIII myosins does not display obvious developmental abnormalities under standard growth conditions (Liu et al. 2026; Talts et al. 2016), complicating genetic dissection of their functions.

In contrast to class XI myosins, for which systematic identification of tail-binding proteins has driven functional understanding, no proteome-wide screen for class VIII myosin-associated proteins has been reported. The few known interacting proteins have each been identified through studies focused on the binding partner or cellular process rather than on the myosin itself. During Agrobacterium-mediated plant transformation, the bacterial effector protein VirE2 binds directly to the cargo-binding domains (CBDs) of ATM2, VIIIA, and VIIIB, and these myosin VIII isoforms tether VirE2 to the plasma membrane to facilitate transformation (Liu et al. 2026). ATM2 also interacts with the PI4P 5-kinase PIP5K2 and promotes PtdIns(4,5)P2 nanodomain formation at the actin–plasma membrane interface in pollen tubes (Heilmann et al. 2025. Preprint). However, these interacting proteins were each discovered through studies centered on the binding partner or cellular process rather than on the myosin itself. No proteome-wide screen for class VIII myosin-associated proteins has been reported, leaving the full repertoire of their associated proteins largely unknown. Given that the systematic identification of tail-binding proteins was instrumental in revealing the diverse functions of class XI myosins, an analogous unbiased approach is essential to uncover the biological significance of class VIII myosins.

A major obstacle to biochemical identification of myosin-associated proteins in plant cells is the destructive nature of cell lysis. Plant cells contain large central vacuoles harboring abundant proteases that are released upon cell disruption, readily degrading interaction partners before they can be captured by co-immunoprecipitation or affinity purification (Plaxton 2019). Additionally, interactions that are transient or weak are easily disrupted during lysis and purification.

Proximity labeling offers a powerful approach that circumvents these limitations. In this method, a promiscuous biotin ligase fused to the protein of interest covalently labels proximal proteins in vivo, enabling their purification under stringent denaturing conditions (Roux et al. 2012). TurboID, an engineered biotin ligase with dramatically enhanced catalytic activity, biotinylates proteins within approximately 10 nm of the bait, capturing both direct interactors and proteins in the same subcellular compartment (Branon et al. 2018). Because biotinylation occurs in living cells before lysis, proximity partners are covalently marked regardless of subsequent proteolytic degradation, and even weak or transient interactions are captured as stable biotin modifications.

In this study, we employed TurboID-based proximity labeling to screen the Arabidopsis proteome for proteins associated with ATM1 without prior assumptions about specific binding partners. We identified 233 non-redundant candidate ATM1-proximal proteins after gene-level consolidation of hits detected in at least two of three independent biological replicates. To prioritize candidates for validation, we combined proximity labeling data with AlphaFold3-based protein complex structure prediction (Abramson et al. 2024). Co-immunoprecipitation confirmed two top-ranked candidates as ATM1-associated proteins: C3H61/AtTZF5 (At5g44260), a tandem zinc finger protein that regulates mRNA stability at processing bodies and stress granules (Pomeranz et al. 2010); and SFH7 (At2g16380), a Sec14-nodulin domain protein that mediates phosphatidic acid transfer from the ER to chloroplasts (Yao et al. 2023). The identification of an RNA-binding protein and a lipid transfer protein as ATM1-associated proteins raises the possibility of previously unrecognized connections between class VIII myosins and post-transcriptional gene regulation or interorganellar lipid transport.

## Results

### Generation of transgenic Arabidopsis plants expressing ATM1 GTD-TurboID

ATM1 is a class VIII myosin consisting of a motor domain, IQ motifs, a coiled-coil region, and a globular tail domain (GTD) that mediates cargo binding (Fig. 1A). To identify proteins that associate with ATM1 in planta, we employed TurboID-based proximity labeling (Branon et al. 2018). TurboID is an engineered biotin ligase that catalyzes rapid biotinylation of proximal proteins within an estimated labeling radius of approximately 10 nm (Fig. 1B). Importantly, because biotinylation occurs in living cells prior to tissue disruption, this approach can capture transient or weak interactions that are prone to disruption during conventional biochemical purification procedures.

**Figure 1.**
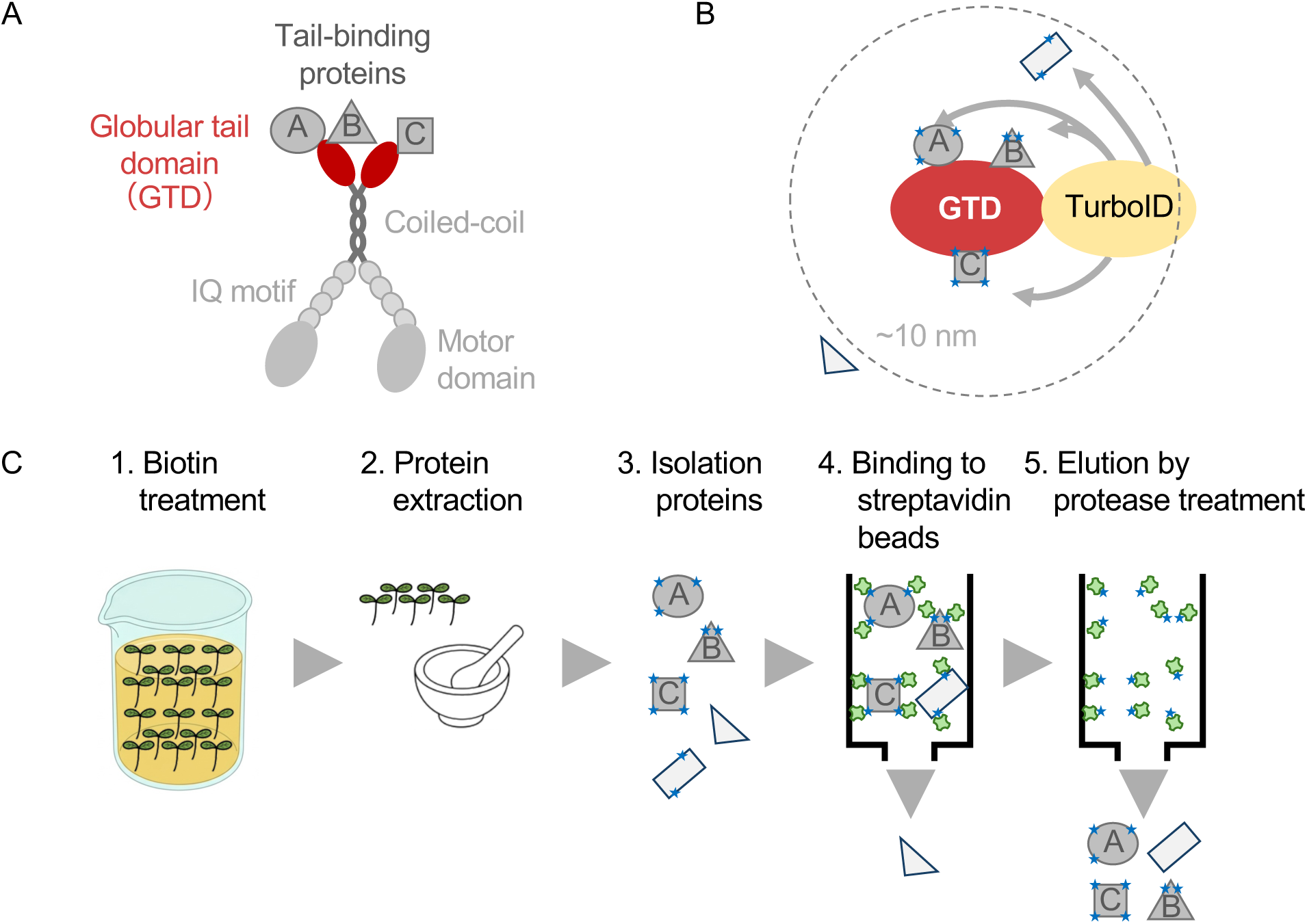
Experimental design for identifying ATM1-proximal proteins by TurboID-based proximity labeling. **(A)** Schematic representation of ATM1 domain structure. ATM1 is a class VIII myosin composed of a motor domain, IQ motifs, a coiled-coil region, and a globular tail domain (GTD, shown in red). The GTD is expected to mediate binding to tail-binding proteins (A, B, C) and contributes to subcellular targeting (Golomb et al. 2008). (B) Principle of TurboID-based proximity labeling. TurboID fused to the ATM1 GTD biotinylates proteins within an approximately 10 nm radius, including direct tail-binding proteins (A, B, C) and other proximal proteins. (C) Experimental workflow. Fourteen-day-old transgenic Arabidopsis seedlings were treated with 500 µM biotin for 3 hours under vacuum infiltration (1). Proteins were extracted by grinding in liquid nitrogen (2), and biotinylated proteins were isolated (3). Biotinylated proteins were captured on streptavidin beads (4) and eluted by on-bead protease digestion for LC-MS/MS analysis (5)

We generated transgenic Arabidopsis thaliana plants expressing either FLAG-ATM1 GTD-TurboID or FLAG-TurboID (control) under the cauliflower mosaic virus 35S promoter. The IQ-tail region of class VIII myosins is sufficient to identify the subcellular structures they are associated with (Golomb et al. 2008). Based on the AlphaFold structural prediction of full-length ATM1, the coiled-coil region of the tail is followed by a structured globular domain corresponding to residues 1031–1166. We therefore used this globular tail domain (GTD) as bait in BioID experiments. Transgene integration was verified by genomic PCR and mRNA expression was confirmed by RT-PCR analysis (Supplementary Fig. S1A, B). Protein expression of the FLAG-tagged fusion proteins was confirmed by immunoblot analysis with anti-FLAG antibody; FLAG-ATM1 GTD-TurboID (∼53 kDa) and FLAG-TurboID (∼38 kDa) were detected in the respective transgenic lines but not in wild-type plants (Supplementary Fig. S1C).

To assess biotinylation activity in vivo, total protein extracts from 14-day-old transgenic seedlings treated with 500 µM biotin for 3 hours were probed with streptavidin-AP. This analysis confirmed biotinylation activity in both transgenic lines (Supplementary Fig. S1D), demonstrating that TurboID was functional in planta.

### Identification of ATM1-proximal proteins by mass spectrometry

Biotinylated proteins were enriched from 14-day-old seedlings using the workflow illustrated in Fig. 1C. Briefly, biotin-treated seedlings were homogenized, and free biotin was removed by gel filtration (PD-10 column). Biotinylated proteins were then purified by streptavidin affinity chromatography and analyzed by liquid chromatography-tandem mass spectrometry (LC-MS/MS). Three independent biological replicates were performed for each genotype.

For each replicate, protein groups identified by DIA-NN (Demichev et al. 2020) were expanded into individual UniProt accessions (The UniProt Consortium, 2023), and redundant sequences (≥90% amino acid similarity) were removed, retaining the entry with the higher UniProt annotation score. This yielded 579, 1,162, and 765 unique proteins for replicates 1, 2, and 3, respectively (Fig. 2A). Of these, 284 proteins were detected in at least two of three replicates, including 35 detected in all three replicates (Fig. 2A). Because the same gene was occasionally represented by different UniProt accessions across replicates, these 284 proteins were consolidated at the TAIR gene locus level (Lamesch et al. 2012), yielding 233 non-redundant candidate ATM1-proximal proteins (Table S1)

**Figure 2.**
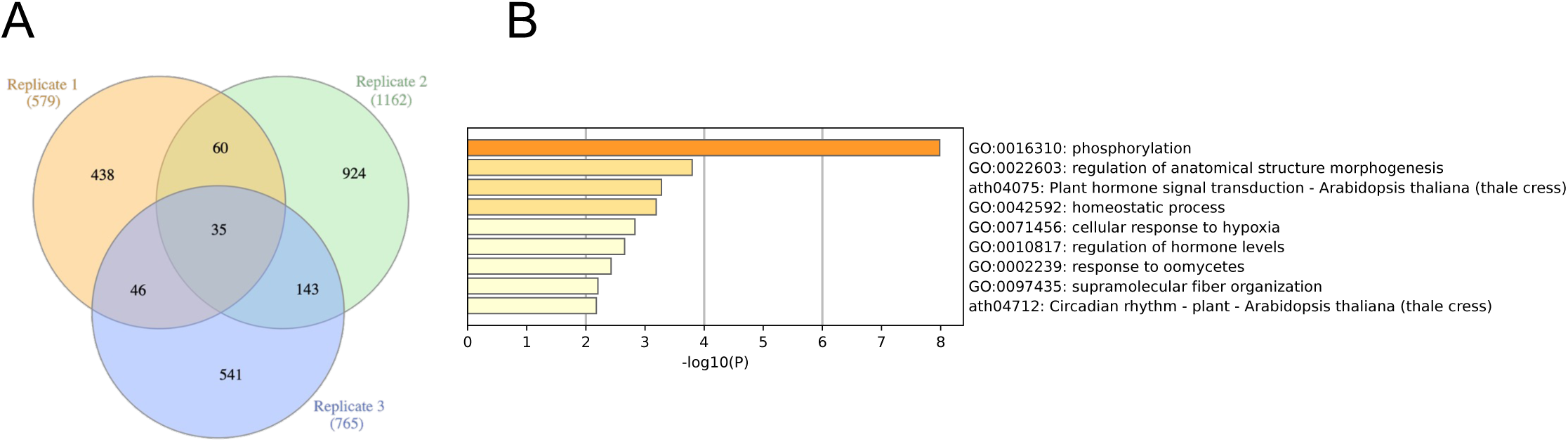
BioID identification and functional enrichment of ATM1-proximal proteins. (A) Venn diagram showing the overlap of proteins identified across three independent biological replicates after sequence redundancy removal. Replicate 1, Replicate 2, and Replicate 3 yielded 579, 1,162, and 765 unique proteins, respectively. A total of 284 proteins were detected in at least two of three replicates; following consolidation to TAIR gene loci, these yielded 233 non-redundant candidate ATM1-proximal proteins. (B) Functional enrichment analysis of the 233 candidate ATM1-proximal proteins was performed using Metascape (Zhou et al. 2019). Of the 233 candidates, 159 genes were mapped to annotated GO terms and used as input; all *Arabidopsis thaliana* genes were used as the background. The bar graph shows representative terms from nine enriched clusters (GO Biological Processes and KEGG Pathways), colored by statistical significance (−log10P). The most significantly enriched term was “phosphorylation” (GO:0016310; 21 genes, P = 1.0 × 10⁻^8^). See Supplementary Table S2 for complete enrichment data.

### Functional enrichment analysis of ATM1-proximal proteins

To characterize the functional composition of the 233 candidate ATM1-proximal proteins, we performed functional enrichment analysis using Metascape (Zhou et al. 2019). Of the 233 candidates, 159 genes could be mapped to annotated GO terms and were used in the analysis. Enriched terms were hierarchically clustered into nine representative groups (Fig. 2B).

The most significantly enriched term was “phosphorylation” (GO:0016310; 21 genes, 13.2% of analyzed candidates; P = 1.0 × 10⁻^8^). The second cluster was “regulation of anatomical structure morphogenesis” (GO:0022603; 8 genes, P = 1.6 × 10⁻^4^). The third cluster comprised KEGG pathways “Plant hormone signal transduction” (ath04075; 12 genes, P = 5.3 × 10⁻^4^) and “Plant–pathogen interaction” (ath04626; 8 genes, P = 7.2 × 10⁻^4^). Additional enriched clusters included “homeostatic process” (GO:0042592; 10 genes, P = 6.5 × 10⁻^4^), “cellular response to hypoxia” (GO:0071456; 8 genes, P = 1.5 × 10⁻^3^), “regulation of hormone levels” (GO:0010817; 8 genes, P = 2.2 × 10⁻^3^), “response to oomycetes” (GO:0002239; 4 genes, P = 3.7 × 10⁻^3^), “supramolecular fiber organization” (GO:0097435; 5 genes, P = 6.3 × 10⁻^3^), and “Circadian rhythm” (ath04712; 3 genes, P = 6.7 × 10⁻^3^) (Fig. 2; Supplementary Table S2).

While functional enrichment analysis characterizes the overall functional landscape of the candidate pool, it does not identify which proteins directly interact with ATM1. We therefore employed computational structure prediction as an orthogonal approach to prioritize candidates for experimental validation.

### AlphaFold3 structure prediction identifies high-confidence interaction candidates

Proximity labeling identifies proteins within the biotinylation radius but does not distinguish direct physical interactors from proteins that are merely located nearby or indirectly associated through multiprotein complexes. In this study, we focused on candidates predicted to form direct complexes with the ATM1 GTD, although the BioID dataset likely also contains functionally relevant indirect interactors. To prioritize candidates most likely to form direct interactions with ATM1, we used AlphaFold3 (Abramson et al. 2024) to predict the complex structure between the ATM1 GTD and each of the 233 candidate proteins.

Predicted complexes were evaluated using two complementary metrics: the interface predicted template modeling (ipTM) score, which assesses overall confidence in the predicted interface, and the interaction prediction score from aligned errors (ipSAE)(Dunbrack Jr 2025), which evaluates the precision of atomic positions at the interface. Three candidates exceeded an ipTM score of 0.6, a threshold below which predictions are generally considered unreliable (Evans et al. 2021). These three candidates were selected for experimental validation by co-immunoprecipitation.

The top 10 candidates ranked by ipTM score are shown in Table 1. C3H61/AtTZF5 (At5g44260) ranked first with ipTM = 0.67 and ipSAE = 0.62. DRB2 (At2g28380) ranked second with ipTM = 0.62 and ipSAE = 0.59. SFH7 (At2g16380) ranked third with ipTM = 0.60 and ipSAE = 0.41 (Table 1).

**Table 1.**
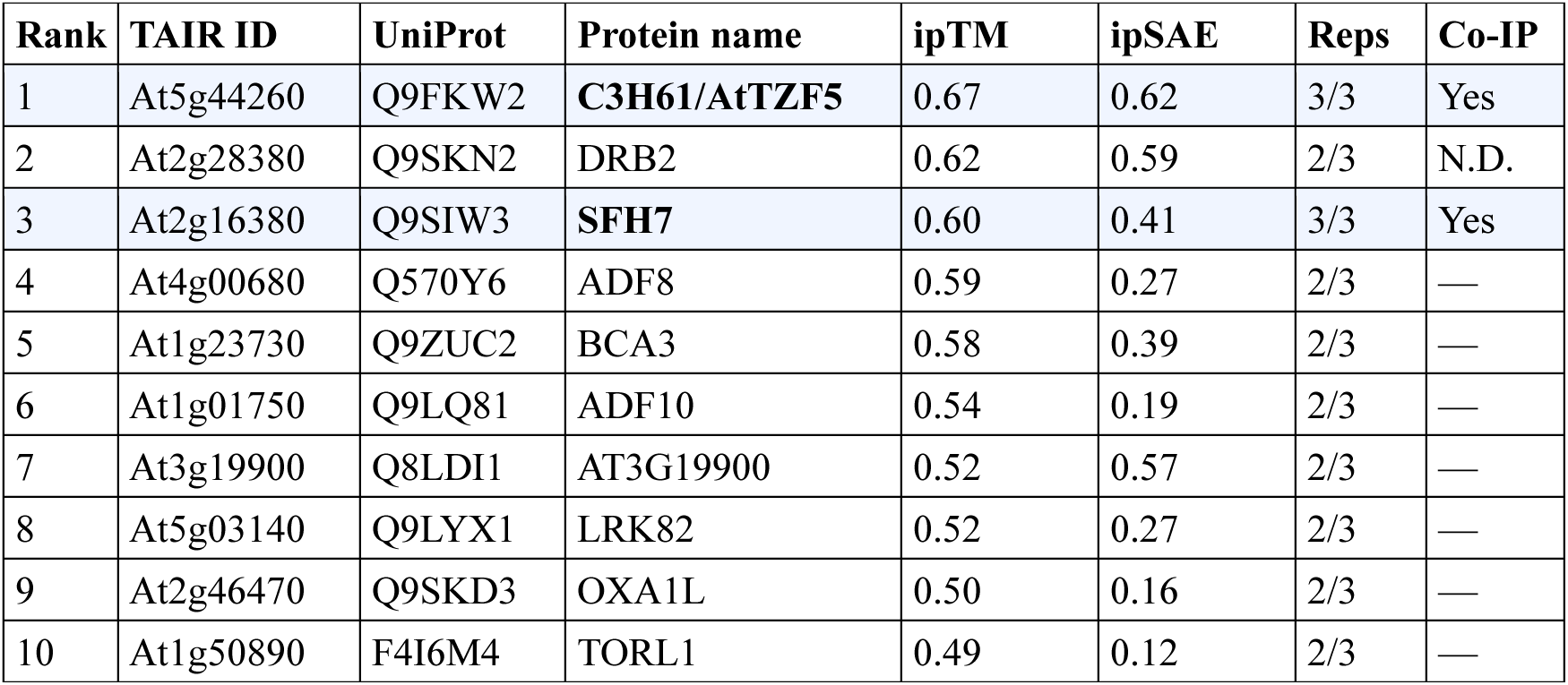
Top 10 candidate ATM1-associated proteins ranked by AlphaFold3 ipTM score. AlphaFold3 was used to predict complex structures between the ATM1 GTD and each of the 233 candidate ATM1-proximal proteins identified by BioID. Predicted complexes were ranked by the interface predicted template modeling score (ipTM), which assesses overall confidence of the predicted interaction interface. The interaction prediction Score from Aligned Errors (ipSAE) (Dunbrack Jr 2025. Preprint) evaluates the precision of predicted atomic positions at the interface. Higher values for both metrics indicate greater confidence. Candidates with ipTM ≥ 0.6 are considered high-confidence predictions (Evans et al. 2021. Preprint). TAIR ID, UniProt accession, and protein names are shown. C3H61 (rank 1) and SFH7 (rank 3) were validated by co-IP (Fig. 4).

The AlphaFold3-predicted structure of the ATM1–C3H61 complex revealed that the zinc-binding motif of C3H61 packs against the ATM1 GTD, forming a well-defined interface (Fig. 3A). C3H61 belongs to the arginine-rich tandem CCCH zinc finger (RR-TZF) protein family, whose members are implicated in RNA binding and mRNA turnover in stress responses and hormone signaling (Bogamuwa and Jang 2014).

**Figure 3.**
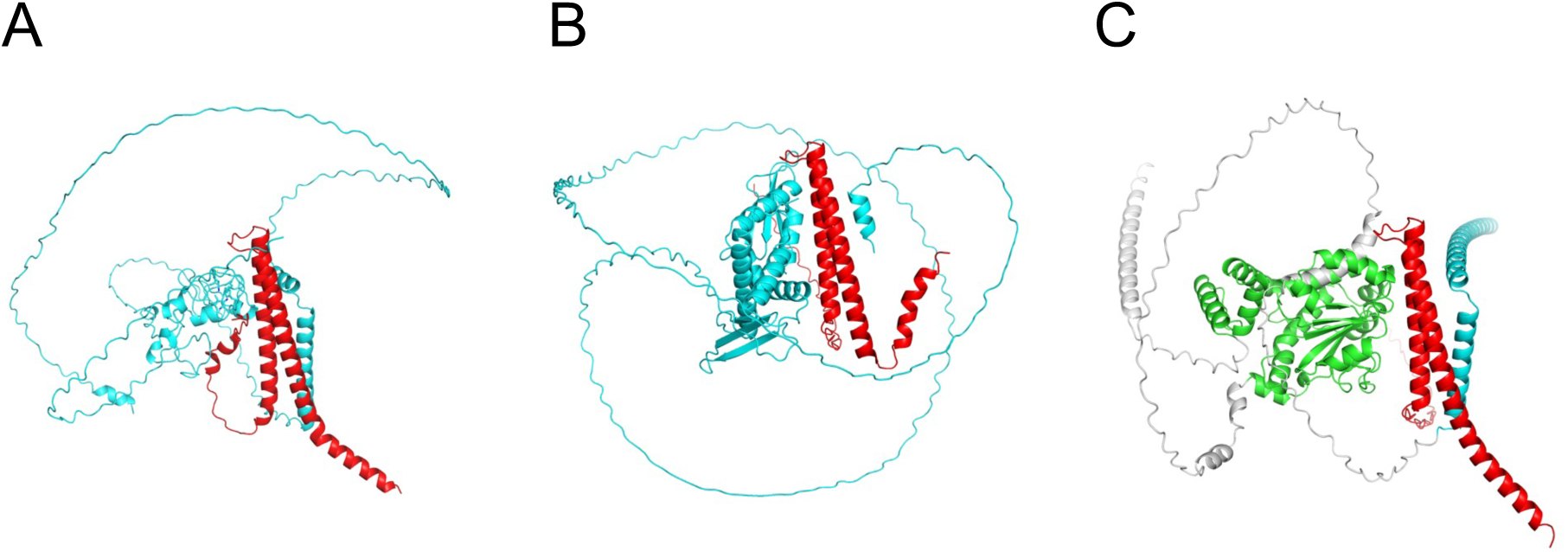
AlphaFold3-predicted complex structures of ATM1 GTD with candidate interaction partners. (A) Predicted structure of the ATM1 GTD–C3H61 complex. The zinc-binding motif of C3H61 (cyan) packs against the ATM1 GTD (red), forming a well-defined interface. C3H61 ranked first among 233 candidates by ipTM score (0.67) and ipSAE score (0.62). (B) Predicted structure of the ATM1 GTD–DRB2 complex. The predicted interface involves the alpha helix at the C-terminal region of DRB2, which comprises the low-confidence (pLDDT) region, indicating structural uncertainty at the predicted interface. DRB2 ranked second by ipTM score (0.62). (C) Predicted structure of the ATM1 GTD–SFH7 complex. The predicted interface involves the Sec14 lipid-binding domain of SFH7 (green), the nodulin domain of SFH7 (cyan), remaining parts of SFH7 (white), and the ATM1 GTD (red). SFH7 ranked third by ipTM score (0.60).

In the predicted ATM1–DRB2 complex, the interaction interface involves the C-terminal region of DRB2 (Fig. 3B). However, the binding area, comprised of two helices of ATM1 and the C-terminal region of DRB2, was small and lacked hydrophilic interactions, implying that the binding may be unstable. Therefore, it was expected that the ipTM between ATM1 and DRB2 would be overestimated due to the existence of the inaccurately predicted α-helix. Furthermore, the C-terminal region showed low per-residue confidence in the AlphaFold3 prediction, indicating structural uncertainty at the predicted interface (Fig. S4A).

The predicted ATM1–SFH7 complex revealed interfaces at which ATM1 is sandwiched by the Sec14 lipid-binding domain and the nodulin domain of SFH7 (Fig. 3C). SFH7 is a Sec14-like phosphatidylinositol transfer protein with an N-terminal Sec14 domain and a C-terminal nodulin domain, and has been shown to mediate phosphatidic acid transport from the endoplasmic reticulum to the chloroplast outer envelope (Yao et al. 2023).

Based on these predictions, C3H61, DRB2, and SFH7—the three candidates with ipTM ≥ 0.6—were selected for experimental validation by co-immunoprecipitation.

### Co-immunoprecipitation validates ATM1 associations with C3H61 and SFH7

To validate the predicted interactions experimentally, we performed co-immunoprecipitation (co-IP) assays in *Nicotiana benthamiana* leaves by transient co-expression of tagged proteins. For these experiments, we used the ATM1 tail domain, comprising the coiled-coil region and the GTD, as bait to better approximate the native conformation of the C-terminal cargo-binding region. GFP-ATM1 tail was immunoprecipitated using anti-GFP magnetic beads, with GFP alone serving as a negative control.

For C3H61 validation, GFP-ATM1 tail or GFP alone was co-expressed with FLAG-C3H61 in *N. benthamiana* leaves (Fig. 4A). Immunoblot analysis of input fractions confirmed expression of all proteins at expected molecular weights: GFP-ATM1 tail at approximately 55 kDa, GFP at approximately 27 kDa, and FLAG-C3H61 as a broad band centered at approximately 44 kDa. The broad migration pattern of FLAG-C3H61 was reproducible across independent experiments and may reflect the arginine-rich intrinsically disordered regions characteristic of the RR-TZF protein family (Bogamuwa and Jang 2014). Direct purification of FLAG-C3H61 using anti-FLAG beads confirmed that the broad migration pattern is an intrinsic property of the C3H61 protein rather than an artifact of indirect pull-down (Supplementary Fig. S2). Following immunoprecipitation with anti-GFP beads, FLAG-C3H61 was detected in the co-IP fraction of GFP-ATM1 tail but not in that of GFP alone (Fig. 4A), demonstrating specific association of C3H61 with the ATM1 tail domain.

**Figure 4.**
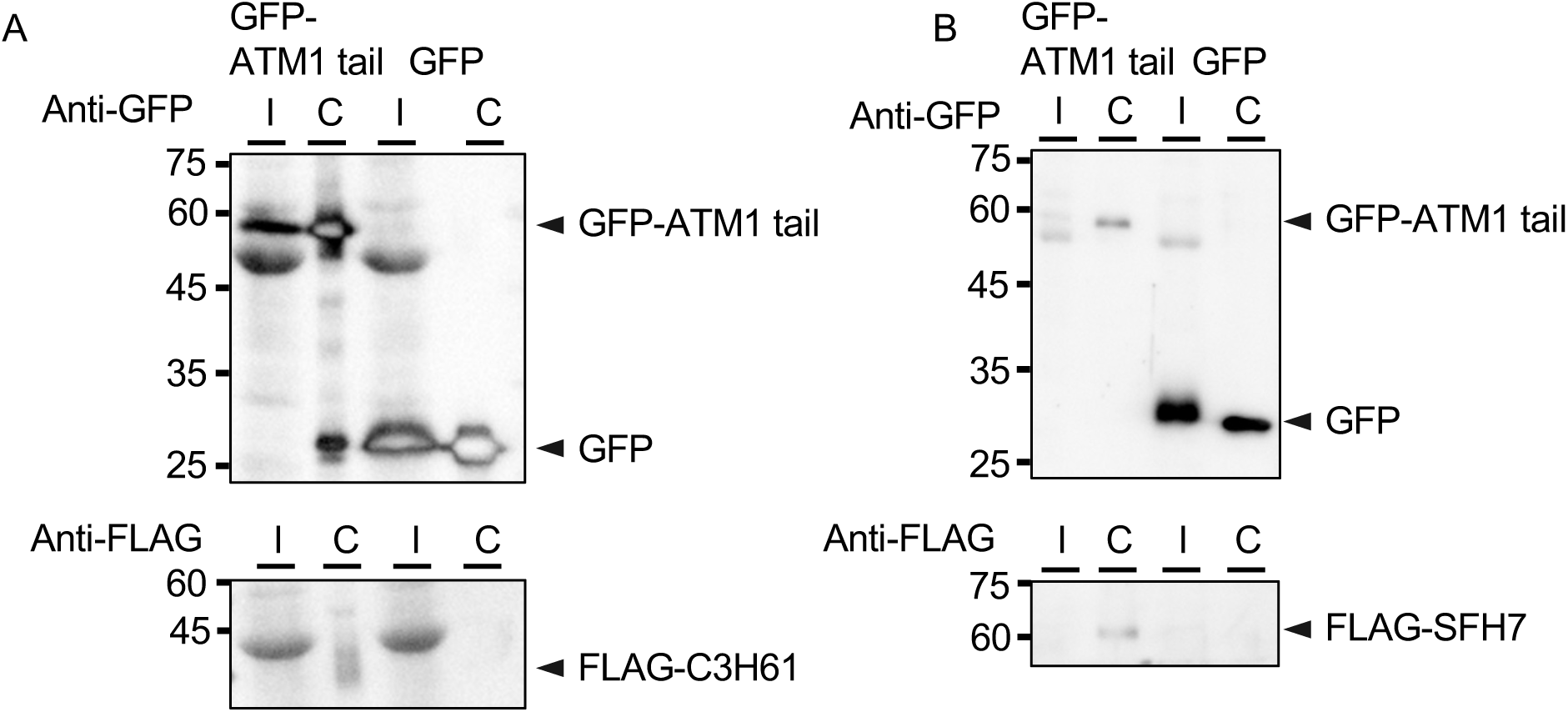
Co-immunoprecipitation validates the association of ATM1 with C3H61 and SFH7. Co-immunoprecipitation (co-IP) assays were performed using *Nicotiana benthamiana* leaves transiently co-expressing the indicated proteins. The ATM1 tail domain (amino acids 930–1166), encompassing both the coiled-coil region and the GTD, was used as bait. GFP-ATM1 tail or GFP alone was co-expressed with FLAG-tagged prey proteins. Protein complexes were immunoprecipitated with anti-GFP magnetic beads. Total protein extracts (I, input) and immunoprecipitated fractions (C, co-IP) were analyzed by immunoblotting with anti-GFP (upper panels) or anti-FLAG (lower panels) antibodies. (A) Co-IP of C3H61 with ATM1 tail. GFP-ATM1 tail (∼55 kDa) or GFP (∼27 kDa) was co-expressed with FLAG-C3H61. Anti-FLAG immunoblotting detected FLAG-C3H61 as a broad band centered at approximately 44 kDa in input fractions. This broad migration pattern may reflect the arginine-rich intrinsically disordered regions characteristic of the RR-TZF protein family (Bogamuwa and Jang, 2014). Immunoprecipitation with anti-FLAG beads confirmed that the broad band corresponds to FLAG-C3H61 (Supplementary Fig. S2). FLAG-C3H61 was co-precipitated with GFP-ATM1 tail but not with GFP alone, demonstrating specific association with the ATM1 tail domain. (B) Co-IP of SFH7 with ATM1 tail. GFP-ATM1 tail (∼55 kDa) or GFP (∼27 kDa) was co-expressed with FLAG-SFH7. Anti-FLAG immunoblotting detected FLAG-SFH7 at approximately 65 kDa. FLAG-SFH7 was co-precipitated with GFP-ATM1 tail but not with GFP alone, demonstrating specific association with the ATM1 tail domain.

For SFH7 validation, the same experimental setup was used with FLAG-SFH7 as prey (Fig. 4B). Input fractions confirmed expression of GFP-ATM1 tail (∼55 kDa), GFP (∼27 kDa), and FLAG-SFH7 (∼65 kDa) at comparable levels in both samples. Following immunoprecipitation with anti-GFP beads, FLAG-SFH7 was detected in the co-IP fraction of GFP-ATM1 tail but not in that of GFP alone (Fig. 4B), demonstrating that SFH7 also specifically associates with the ATM1 tail domain.

DRB2, the second-ranked candidate (ipTM = 0.62), was also subjected to co-IP analysis; however, insufficient expression of FLAG-DRB2 in *N. benthamiana* leaves precluded reliable assessment of the interaction (Supplementary Fig. S3).

Taken together, these results identify C3H61 and SFH7 as proteins that associate with the ATM1 tail domain in co-immunoprecipitation assays. Both proteins were detected in multiple BioID replicates, showed ipTM scores above the 0.6 confidence threshold, and were confirmed by co-IP with GFP-alone negative controls.

## Discussion

We employed TurboID-based proximity labeling to identify proteins associated with the globular tail domain (GTD) of ATM1, a class VIII myosin in Arabidopsis. This approach identified 233 non-redundant candidate ATM1-proximal proteins after gene-level consolidation of hits detected in at least two of three independent biological replicates, from which we selected candidates for validation using a combination of AlphaFold3-based structure prediction and co-IP. We confirmed two proteins as ATM1-associated proteins: C3H61/AtTZF5, a tandem zinc finger protein involved in post-transcriptional gene regulation, and SFH7, a phosphatidic acid transfer protein. These represent the first identified interaction partners of ATM1 and suggest that this class VIII myosin may be associated with proteins involved in post-transcriptional gene regulation and lipid transfer.

### Combining Proximity Labeling with AlphaFold3 Structure Prediction

TurboID biotinylates proteins in close proximity to the bait protein in vivo, enabling detection of weak or transient interactions that may be lost during conventional affinity purification (Branon et al. 2018). However, this sensitivity comes at the cost of specificity: proteins may be biotinylated simply due to spatial proximity without direct physical contact with the bait. To address this limitation, we employed AlphaFold3 to predict potential binding interfaces between ATM1 and the candidate proteins (Abramson et al. 2024). AlphaFold3 predicts direct physical contacts between protein residues; candidates with high ipTM scores are therefore predicted to form direct binding interfaces with ATM1. By ranking candidates based on ipTM scores, we could prioritize proteins most likely to interact directly with ATM1 for experimental validation.

Co-IP validation was successful for C3H61 and SFH7, both of which showed predicted interfaces at well-folded structural domains (the zinc finger motif and the Sec14 lipid-binding domain and nodulin domain, respectively). By contrast, the predicted DRB2 interface was small and lacked hydrophilic interactions, and the C-terminal region exhibited low pLDDT scores (Supplementary Fig. S4A), suggesting that the ipTM score of 0.62 should be interpreted with caution. Furthermore, co-IP validation of DRB2 was not possible due to insufficient FLAG-DRB2 expression in *N. benthamiana* (Supplementary Fig. S3), and alternative experimental approaches will be needed to assess this candidate.

These observations suggest that combining multiple criteria—high ipTM scores, well-defined structural interfaces, and high BioID reproducibility—may improve the success rate of candidate validation. The validation rate in our study (two of two assessable candidates confirmed by co-IP) is consistent with the utility of this approach.

### ATM1 Associates with an RNA-Binding Protein

The identification of C3H61/AtTZF5 as an ATM1-associated protein provides the first link between a class VIII myosin and an RNA-binding protein. C3H61 ranked first among all 233 candidates in our AlphaFold3 analysis (ipTM = 0.67), was detected in all three BioID replicates, and was confirmed by co-IP. Immunoprecipitation with anti-FLAG beads further verified that the co-precipitated broad band corresponds to FLAG-C3H61.

C3H61/AtTZF5 belongs to the plant-specific RR-TZF (arginine-rich tandem CCCH zinc finger) protein family, which comprises 11 members in Arabidopsis (Wang et al. 2008). These proteins are characterized by two tandem zinc finger motifs preceded by an arginine-rich region, a domain architecture that enables RNA binding (Bogamuwa and Jang 2014). AtTZF proteins localize to processing bodies (P-bodies) and stress granules, cytoplasmic ribonucleoprotein condensates where mRNA decay and translational repression occur (Bogamuwa and Jang 2016; Pomeranz et al. 2010). Notably, class XI myosin XI-K has been reported to associate with DCP1, a P-body component (Steffens et al. 2014), suggesting that myosin–RNA granule protein connections may be a broader theme in plant biology.

The physiological role of the ATM1–C3H61 association and the subcellular site where it occurs remain to be elucidated. Future studies examining whether atm1 mutants exhibit altered P-body dynamics or mRNA turnover will be important for understanding the functional significance of this association.

### ATM1 Associates with a Lipid Transfer Protein

The identification of SFH7 as an ATM1-associated protein provides the first link between a class VIII myosin and a lipid transfer protein. SFH7 belongs to the Sec14-nodulin domain protein family and has recently been characterized as a phosphatidic acid (PA) transfer protein (Yao et al. 2023). SFH7 was detected in all three BioID replicates. Together with its paralog SFH5, SFH7 mediates the non-vesicular transfer of PA from the endoplasmic reticulum to the chloroplast outer envelope membrane, where PA serves as a precursor for the synthesis of galactolipids including monogalactosyldiacylglycerol (MGDG). The sfh5 sfh7 double mutant exhibits defective chloroplast development, reduced MGDG content, and abnormal thylakoid membrane organization (Yao et al. 2023).

AlphaFold3 predicted the potential binding of SFH7 to be the Sec14-like lipid-binding domain and the nodulin domain (Fig. 3C). The interface between the Sec14-like domain and SFH7 included a low confidence region (pLDDT < 0.5) (Fig. S4B) and contained only one salt bridge between D1090 (ATM1) and K216 (SFH7). On the other hand, the three-helix bundle formed by ATM1-GTD and SFH7 nodulin domain (binding around residues 464-500) had both a hydrophilic patch and salt bridges (K1117-D490, K1124-D485, E1082-K480, and K1135-E478, where the former residue corresponds to ATM1 and the latter to SFH7, respectively), presenting a more plausible interface.

The association was confirmed by co-IP: FLAG-SFH7 was specifically co-precipitated with GFP-ATM1 tail but not with GFP alone. Mutating the residues listed above may provide a more detailed mechanism of the ATM1-SFH7 binding; however, such an analysis is beyond the scope of this paper.

SFH5 and SFH7 are both expressed in seedlings and colocalize at the ER (Yao et al. 2023), yet SFH5 was not detected in any BioID replicate. This specificity suggests that the ATM1–SFH7 association is mediated by structural features unique to SFH7, rather than reflecting a general proximity to PA transfer proteins.

The physiological role of the ATM1–SFH7 association and the subcellular site where it occurs remain to be elucidated. Future studies examining whether atm1 mutants exhibit defects in chloroplast lipid composition or ER–chloroplast contact site organization will help clarify the functional significance of this association. It should also be noted that the 233 candidate proteins identified by proximity labeling likely include indirect interactors associated through multiprotein complexes, which were beyond the scope of our pairwise AlphaFold3 predictions; exploring these indirect associations will be an important future direction.

## Conclusions

In summary, we have used TurboID-based proximity labeling combined with AlphaFold3 structure prediction to identify candidate ATM1-associated proteins, and confirmed the association of C3H61/AtTZF5 and SFH7 by co-IP. Both validated proteins were detected in all three BioID replicates and showed well-defined predicted direct binding interfaces with structured domains.

These findings demonstrate that a systematic, unbiased identification of tail-binding proteins—the approach that proved instrumental in elucidating class XI myosin functions—can similarly reveal previously unrecognized cellular roles of class VIII myosins, even where genetic approaches have been limited by functional redundancy. More broadly, the workflow employed here—comprehensive collection of proximal protein candidates by proximity labeling in vivo, followed by computational prioritization through AlphaFold3 complex structure prediction, and targeted experimental validation of high-confidence candidates—provides a generally applicable strategy for identifying protein–protein interactions in any eukaryotic system. This approach is particularly advantageous for proteins whose interaction partners are unknown and for which no prior assumptions can guide candidate selection. Our results provide the first insights into the protein interaction network of a class VIII myosin, linking ATM1 to post-transcriptional gene regulation and interorganellar lipid transport—two cellular functions not previously connected to class VIII myosins.

## Materials and Methods

### Vector construction

Vectors were constructed using Gateway Technology (Invitrogen, Carlsbad, CA, USA) with destination vectors pH2GW7 (Karimi et al 2002), pGWB405, pGWB406 and pGWB612 (Nakagawa et al. 2007).

For BioID proximity labeling in Arabidopsis:

ATM1 full-length was PCR amplified from pENTR ATM1 Full (provided by M. Tominaga, Waseda University) using forward primer (5′-ACAAAAAAGCAGGCTCTATGGCTCAGAAGGTTACTC-3′) and reverse primer (5′-ACAAGAAAGCTGGGTTCAATACCTGGTGCTATTTCTCC-3′), and cloned into linearized pDONR/Zeo using In-Fusion cloning to generate pDONR/Zeo ATM1 Full-length.

To create the TurboID fusion construct, pDONR/Zeo ATM1 Full-length was linearized using forward primer (5′-ACCCAGCTTTCTTGTACAAAGTTG-3′) and reverse primer (5′-ATACCTGGTGCTATTTCTCCTTCC-3′). TurboID with a GSGSGS linker was PCR amplified from pENTR221 V5turboID-GFP (provided by M. Nakamura, Saitama University) using forward primer (5′-GGAGAAATAGCACCAGGTATGGCGGCAGCGGCTCTGGAAAAGACAATACTGTGCCTCTG-3′) and reverse primer (5′-TTTGTACAAGAAAGCTGGGTTCACTTTTCGGCAGACCGCA-3′).

The TurboID fragment was inserted into linearized ATM1 full-length using In-Fusion cloning to generate pDONR/Zeo ATM1 Full-length TurboID.

To obtain ATM1 GTD (amino acids 1031–1166) fused to TurboID, the sequence encoding the motor domain, IQ motifs, and coiled-coil region (amino acids 1–1030) was deleted from pDONR/Zeo ATM1 Full-length TurboID using the PrimeSTAR Mutagenesis system (Takara). Outward-facing primers were designed to amplify the entire plasmid excluding the HMM-encoding region: forward primer (5′-AGGCTCTAGAAACTCAGATGCATCT-3′) and reverse primer (5′-GAGTTTCTAGAGCCTGCTTTTTTGTA-3′). The resulting PCR product self-circularized and was transformed into E. coli, yielding pDONR/Zeo ATM1 GTD TurboID.

As a control, TurboID alone was PCR amplified from pENTR221 V5turboID-GFP using forward primer (5′-ACAAAAAAGCAGGCTCTAAAGACAATACTGTGCCTCTG-3′) and reverse primer (5′-TACAAGAAAGCTGGGTTCCAGAGCCGCTGCCGCCCTTTTCGGCAGACCGCA-3′), cloned into pDONR/Zeo, and transferred to pGWB612 by LR recombination to generate 35S:FLAG-ATM1 GTD-TurboID and 35S:FLAG-TurboID expression vectors.

For co-immunoprecipitation in *N. benthamiana*:

ATM1 tail (amino acids 930–1166) was PCR amplified using forward primer (5′-TTTGTACAAAAAAGCAGGCTCTTGTTCAGGGGATATGGATG-3′) and reverse primer (5′-TTTGTACAAGAAAGCTGGGTTCAATACCTGGTGCTATTTCTCC-3′), cloned into pDONR/Zeo, and transferred to pGWB406 to generate 35S:GFP-ATM1tail. As a GFP control construct, pH2GW7-GFP was generated as follows. The *GFP* gene was PCR amplified from pGWB405 using forward primer (5′-gcaggctccgcggccATGgtgagcaagggcgagga-3′) and reverse primer (5′-agctgggtcggcgcgTTActtgtacagctcgtcca-3′). The entry clone pENTR-GFP was produced by ligating the *GFP* PCR product into pENTR-CUT (Yamaguchi et al. 2025) using an In-Fusion Cloning Kit (Takara). pENTR-GFP was recombined into the destination vector pH2GW7 via LR Clonase II (Invitrogen), creating the expression vector pH2GW7-GFP.

C3H61 (At5g44260), SFH7 (At2g16380), and DRB2 (At2g28380) were each amplified from *A. thaliana* cDNA by nested PCR. First-round primers were: for C3H61, forward (5′-CTCTTCCTATCTCTCTCTCATTCAAACCC-3′) and reverse (5′-CGCCGTATAATTATTCCCGTCG-3′); for SFH7, forward (5′-CATCTCCAATAAGGTGTTTCATCAAAAGTC-3′) and reverse (5′-CTTTTCATTTGATATTTTCGCGGG-3′); for DRB2, forward (5′-CCATGATTCGAAGATACAGAGCCT-3′) and reverse (5′-CTCCCCAATATAATACAAATCTTTCGG-3′). Second-round primers containing In-Fusion adaptor sequences were: for C3H61, forward (5′-TTTGTACAAAAAAGCAGGCTCTATGGACGTCGAACATCAC-3′) and reverse (5′-TTTGTACAAGAAAGCTGGGTTCATGTCAAAAGATCGTTCA-3′); for SFH7, forward (5′-TTTGTACAAAAAAGCAGGCTCTATGGCTGACACCAAACA-3′) and reverse (5′-TTTGTACAAGAAAGCTGGGTTAGAACCTGAAGAACATTCTC-3′); for DRB2, forward (5′-TTTGTACAAAAAAGCAGGCTCTATGTATAAGAACCAGCTACAAG-3′) and reverse (5′-TTTGTACAAGAAAGCTGGGTTCAGATCTTTAGGTTCTCCA-3′). Each PCR product was cloned into pDONR/Zeo and transferred to pGWB612 to generate 35S:FLAG-C3H61, 35S:FLAG-SFH7, and 35S:FLAG-DRB2.

### Plant materials and growth conditions

*Arabidopsis thaliana* Columbia-0 (Col-0) ecotype was used throughout this study. Transgenic lines were generated by Agrobacterium-mediated floral dip transformation. Seeds were surface-sterilized with 70% ethanol and stratified at 4°C for 48 hours. Seedlings were grown on half-strength Murashige and Skoog medium supplemented with 0.5% sucrose under long-day conditions (16 h light / 8 h dark) at 22°C with a photon flux density of 100 μmol m^−2^ s^−1^.

*Nicotiana benthamiana* plants were grown in soil under the same photoperiod conditions at 25°C for 4–6 weeks before use.

### Proximity labeling treatment

14-day-old seedlings (approximately 0.5–1.0 g fresh weight) were harvested and submerged in 500 μM D-biotin in half-strength MS liquid medium. Vacuum infiltration was applied for 15 minutes, followed by incubation at room temperature for 3 hours. Seedlings were washed six times with approximately 200 mL ice-cold water to remove excess biotin, dried on absorbent paper, and immediately flash-frozen in liquid nitrogen. Frozen tissue was ground to a fine powder using a mortar and pestle in liquid nitrogen. Control samples expressing FLAG-TurboID alone were processed identically.

### Protein extraction and sample preparation

Frozen plant powder was resuspended in extraction buffer (50 mM Tris-HCl pH 7.5, 150 mM NaCl, 0.1% SDS, 1% Triton X-100, 1 mM EGTA, 1 mM DTT, and protease inhibitor cocktail) and rotated at 4°C for 10 minutes. Lysates were sonicated with an SND US-2 (four 30-second pulses with 1.5-minute intervals on ice) and clarified by sequential centrifugation at 3,000 × g for 5 minutes and 15,000 × g for 15 minutes at 4°C.

### Removal of free biotin and affinity enrichment of biotinylated proteins

Free biotin was removed by gel filtration on PD-10 desalting columns (GE Healthcare) equilibrated with PBS containing 1 mM DTT. The desalted extract was incubated with 50 μL of μMACS Streptavidin MicroBeads (Miltenyi Biotec) for 3 hours at 4°C. Beads were captured on μMACS columns (Miltenyi Biotec) and washed ten times with 200 μL PBS containing 1 mM DTT.

### On-column proteolytic digestion

Columns were washed with 50 mM Tris-HCl pH 7.5 to remove detergents. Proteins were digested on-column using LysC (5 ng/μL) and trypsin (5 ng/μL) in 2 M urea, 50 mM Tris-HCl pH 7.5, 1 mM DTT for 30 minutes at 26°C, followed by alkylation with 5 mM chloroacetamide in the same buffer for 16–20 hours at 26°C. Digested samples were stored at −80°C until analysis.

### LC-MS/MS analysis and data processing

Peptide mixtures were analyzed by nanoLC–MS/MS on a Q Exactive Orbitrap mass spectrometer (Thermo Fisher Scientific, Waltham, MA, USA) coupled to an UltiMate 3000 nanoLC system (Thermo Fisher Scientific) via a nano-electrospray ion source (AMR Inc., Tokyo, Japan). Data-independent acquisition (DIA) was performed with peptide separation on a C18 reversed-phase column using an acetonitrile gradient in 0.1% acetic acid.

Peptide identification and label-free quantification were performed with DIA-NN1.8.1 (Demichev et al. 2020) against the *Arabidopsis thaliana*TAIR10 protein database. Search parameters included trypsin as the digestion enzyme with up to two missed cleavages, carbamidomethylation of cysteine as a fixed modification, and oxidation of methionine and protein N-terminal acetylation as variable modifications. Identifications were filtered at a false discovery rate of 1%. Proteins not detected in any ATM1 GTD-TurboID sample were excluded; missing values in detected samples were imputed with log_2_(4). The mass spectrometry data have been deposited to jPOST (Okuda et al. 2025).

### Candidate protein selection pipeline

For each biological replicate, protein groups reported by DIA-NN were expanded by listing all constituent UniProt accession IDs individually. Amino acid sequences were retrieved from UniProt, and sequence redundancy was removed at a 90% similarity threshold using the entry with the higher UniProt annotation score as the representative. This yielded 579, 1,162, and 765 unique proteins for replicates 1, 2, and 3, respectively.

Proteins detected in at least two of three replicates (284 proteins at the UniProt accession level) were mapped to TAIR gene loci to resolve cases where the same gene was represented by different UniProt accessions across replicates. Representative UniProt accessions for each locus were selected based on protein existence level, annotation score, and sequence length, yielding 233 non-redundant candidate ATM1-proximal proteins (Supplementary Table S1).

### Gene Ontology enrichment analysis

Gene Ontology (GO) enrichment analysis was performed on the 233 candidate ATM1-proximal proteins using Metascape (Zhou et al. 2019) with default settings. Of the 233 candidates, 159 genes were mapped to annotated GO terms and used as input; the complete *Arabidopsis thaliana* proteome served as the background. Enriched terms were hierarchically clustered based on Kappa-statistical similarity, and representative terms from each cluster are reported. Protein–protein interaction (PPI) network analysis and MCODE-based module detection were performed within Metascape using the same input gene list.

### AlphaFold3 structure prediction

AlphaFold3 (Abramson et al. 2024) was used to predict complex structures between the ATM1 GTD (residues 1031–1166) and each of the 233 candidate proteins. We used the local version of AlphaFold3 (commit hash 344735c3) with default parameters. PDB files for templates were retrieved on 15th May 2025. For each candidate complex, 5 (default) models were predicted with different initial pseudorandom number seeds, and the structure with the best AlphaFold 3’s ranking score (weighted sum of the ipTM score, the monomer prediction score, and structural features) was used in the subsequent analyses.

Predicted complexes were evaluated using the interface predicted template modeling score (ipTM) and the interaction prediction Score from Aligned Errors (ipSAE) (Dunbrack Jr 2025. Preprint) using the distance cutoff parameter of 15 Å. Candidates with ipTM ≥ 0.6 were considered high-confidence interaction predictions (Evans et al. 2021. Preprint). Structures were visualized by PyMOL (The PyMOL Molecular Graphics System, Version 2.5.0, Schrödinger, LLC.).

### Transient expression in *N. benthamiana* leaves

Agrobacterium tumefaciens strain GV3101 harboring expression constructs was cultured in LB medium (100 μg/mL spectinomycin, 50 μg/mL rifampicin) at 28°C for 2 days. Bacterial pellets were resuspended in infiltration buffer (10 mM MES-KOH pH 5.7, 10 mM MgCl_2_, 150 μM acetosyringone) to OD600 = 0.3–0.5. For co-expression, suspensions carrying different constructs and the p19 silencing suppressor were mixed at equal volumes. The mixture was infiltrated into the abaxial surface of four-week-old *N. benthamiana* leaves using a needleless syringe. Infiltrated plants were maintained at 25°C (16 h light / 8 h dark) for 48 hours before harvest.

### Co-immunoprecipitation

Approximately 1 g of infiltrated leaf tissue was ground on ice in 1.5–2.0 mL lysis buffer (150 mM NaCl, 1% Ecosurf EH-9, 50 mM Tris-HCl pH 8.0, protease inhibitors: 5 μg/mL leupeptin, 50 μg/mL TLCK, 0.1 mM PMSF, 0.8% protease inhibitor cocktail, 10 μg/mL chymostatin/pepstatin, 40 μg/mL TATP, 1 mM DTT). The homogenate was filtered through a 70 μm PluriStrainer and clarified by centrifugation at 1,000 × g for 5 minutes followed by 15,000 × g for 1 minute at 4°C.

Clarified extract (950 μL) was incubated with 50 μL anti-GFP MicroBeads (Miltenyi Biotec) on ice for 30 minutes. An aliquot of 120 μL was reserved as input. Immunoprecipitation was performed on μMACS columns equilibrated with lysis buffer. Columns were washed five times with 200 μL lysis buffer and once with 200 μL wash buffer (20 mM Tris-HCl pH 7.5). Bound proteins were eluted with pre-heated (95°C) elution buffer (50 mM Tris-HCl pH 6.8, 50 mM DTT, 1% SDS, 1 mM EDTA, 0.005% bromophenol blue, 10% glycerol). For reverse co-IP, anti-FLAG MicroBeads were used in place of anti-GFP MicroBeads with the same protocol.

### Western blot analysis

Protein samples were separated on 10% SDS-PAGE gels and transferred to PVDF membranes (Millipore) at 400 V, 200 mA for 60 minutes in transfer buffer (25 mM Tris-HCl pH 8.0, 192 mM glycine, 20% methanol). Membranes were blocked with 5% skim milk in TBS-T (25 mM Tris-HCl pH 7.5, 150 mM NaCl, 0.05% Tween-20) for 1 hour at room temperature.

For detection of FLAG-tagged proteins, membranes were incubated overnight at 4°C with anti-DYKDDDDK tag monoclonal antibody (clone 1E6; FUJIFILM Wako, Cat# 014-22383) at 1:2,000 in 5% skim milk / TBS-T. For detection of GFP-tagged proteins, anti-GFP antibody (JL-8; Clontech) was used at 1:2,000 in 5% skim milk / TBS-T. After three washes with TBS-T, membranes were incubated with HRP-conjugated anti-mouse IgG secondary antibody (NA931VS; GE Healthcare) at 1:5,000 in TBS-T for 1 hour at room temperature. Signals were detected using an ECL chemiluminescent system and imaged with an iBright imaging system (Thermo Fisher Scientific).

### AlphaFold3 structure prediction — ranking metrics

For each predicted ATM1–candidate complex, two interface quality metrics were calculated: (1) the interface predicted template modeling score (ipTM), which reflects global confidence in the predicted interaction; and (2) the interface surface area error (Dunbrack Jr 2025), which focuses on high-confidence interface residue pairs filtered by predicted aligned error. Candidates were ranked by ipTM in descending order.

### Statistical analysis

Quantitative data are presented as mean ± standard deviation (SD). The number of biological replicates is indicated in figure legends.

## Supporting information

Supplementary Table S1: Complete list of 233 candidate ATM1-proximal proteins

Supplementary Table S2: Complete functional enrichment analysis results

## Accession numbers

Sequence data from this article can be found in TAIR under the following accession numbers: ATM1 (At3g19960), C3H61/AtTZF5 (At5g44260), SFH7 (At2g16380), DRB2 (At2g28380).

## Author Contributions

T.H., K.I., M.T., K.T., and J.O. designed research; T.H., S.N., Y.O., S.S., E.M.-S., and T.L.S. performed research; S.N., S.S., E.M.-S., T.H., and K.I. analyzed data; and T.H. and K.I. wrote the paper.

## Acknowledgements

We thank Prof. Tsuyoshi Nakagawa (Shimane University) for providing the pGWB destination vectors, and Dr. Masayoshi Nakamura (Saitama University) for providing the pENTR221 V5-TurboID-GFP vector.

## Abbreviations

AP: alkaline phosphatase
BioID: proximity-dependent biotin identification
CaMV: cauliflower mosaic virus
Co-IP: co-immunoprecipitation
DIA-NN: Data-Independent Acquisition by Neural Networks
ER: endoplasmic reticulum
GFP: green fluorescent protein
GO: Gene Ontology
GTD: globular tail domain
IDR: intrinsically disordered region
ipSAE: interaction prediction Score from Aligned Errors
ipTM: interface predicted Template Modeling score
KEGG: Kyoto Encyclopedia of Genes and Genomes
MS: mass spectrometry
PA: phosphatidic acid
PD: plasmodesmata
RT-PCR: reverse transcription polymerase chain reaction
SDS-PAGE: sodium dodecyl sulfate-polyacrylamide gel electrophoresis
TAIR: The Arabidopsis Information Resource

## Supplementary Methods

### Genomic DNA extraction

A small leaf (approximately the inner diameter of a 1.5 mL tube lid) was collected from each transgenic line. The leaf tissue was placed in a 1.5 mL tube with a steel bead and frozen at −80°C overnight. Tissue was disrupted using a Tissue Lyser II (Qiagen) at 20/s for 1 minute. Four hundred microliters of extraction buffer (200 mM Tris-HCl pH 7.5, 250 mM NaCl, 25 mM EDTA, 0.5% SDS) was added, and the sample was vortexed briefly. After centrifugation at 14,000 rpm for 1 minute at room temperature, 300 μL of supernatant was transferred to a new tube. DNA was precipitated by addition of 300 μL isopropanol, incubated for 2 minutes at room temperature, and pelleted at 14,000 rpm for 5 minutes. The pellet was air-dried for 10–15 minutes and dissolved in 100 μL TE buffer. The DNA extract was used at 5–50% (v/v) as template in subsequent PCR reactions.

### Transgene verification by genomic PCR

Genomic PCR was performed using Mighty Amp DNA Polymerase Ver.3 (Takara) according to the manufacturer’s instructions. The following primer pairs and expected amplicon sizes were used: for 35S:FLAG-ATM1 GTD-TurboID, 5′-GGAGAAATAGCACCAGGTATG-3′ and 5′-CACAGGAAACAGCTATGACC-3′ (expected product 1,452 bp); for 35S:FLAG-TurboID, 5′-CAATCCCACTATCCTTCG-3′ and 5′-CACAGGAAACAGCTATGACC-3′ (expected product 1,574 bp). Wild-type plants and Milli-Q water were used as negative controls.

### RNA extraction

Total RNA was extracted from 14-day-old seedlings using the Maxwell RSC Plant RNA Kit (Promega, Cat# AS1500) with the Maxwell RSC automated nucleic acid purification instrument, according to the manufacturer’s instructions. On-column DNase I treatment was included to remove genomic DNA contamination. RNA was eluted in 50 μL nuclease-free water.

### Reverse transcription

First-strand cDNA was synthesized from 500 ng total RNA using the PrimeScript RT Reagent Kit with gDNA Eraser (Takara), according to the manufacturer’s instructions. The gDNA elimination reaction was performed at 42°C for 2 minutes prior to reverse transcription.

### RT-PCR

RT-PCR was performed using cDNA as template. For detection of the 35S:FLAG-ATM1 GTD-TurboID transcript, primers 5′-GGAGAAATAGCACCAGGTATG-3′ and 5′-ATCCAGGTTTGATAACTCCGTC-3′ were used (expected product 961 bp from cDNA). As a positive control for cDNA quality, ACTIN2 was amplified using 5′-AGAGATTCAGATGCCCAGAAGTCTTGTTCC-3′ and 5′-AACGATTCCTGGACCTGCCTCATCATACTC-3′ (expected products 353 bp from cDNA and 439 bp from genomic DNA). Genomic DNA from wild-type plants was included as a size control to confirm cDNA-specific amplification, and Milli-Q water was used as a no-template negative control. PCR products were separated on 1% agarose gels and visualized by ethidium bromide staining.

### Streptavidin-AP western blot for in vivo biotinylation

Protein samples from biotin-treated seedlings were extracted in Flag buffer (25 mM HEPES pH 7.4, 150 mM KCl, 4 mM MgCl_2_, 1 mM EGTA, 10% glycerol, 10 mM DTT, and protease inhibitors) by grinding in liquid nitrogen followed by centrifugation at 14,000 rpm for 5 minutes at 4°C. Samples were boiled in SDS sample buffer and separated on 10% SDS-PAGE gels. Proteins were transferred to PVDF membranes at 400 V, 200 mA for 60 minutes.

Membranes were washed three times with TBS-T (2, 3, and 5 minutes) and blocked overnight at 25°C in 0.1% gelatin / TBS-T. Membranes were then incubated with Streptavidin-AP (V559C; Promega) diluted 1:1,000 in 0.1% gelatin / TBS-T for 1 hour at 25°C. After three washes with TBS-T (2, 3, and 5 minutes), biotinylated proteins were visualized by alkaline phosphatase (AP) colorimetric detection using NBT (20 μL of stock per 5 mL AP buffer) and BCIP (10 μL of stock per 5 mL AP buffer). The reaction was stopped by rinsing with water, and membranes were scanned.

## Supplementary Tables

**Supplementary Table S1. Complete list of 233 candidate ATM1-proximal proteins.** List of all 233 non-redundant proteins detected in at least two of three BioID replicates after TAIR gene locus consolidation. For each protein, the following information is shown: TAIR ID, representative UniProt accession, gene name, protein length (amino acids), AlphaFold3 ipTM score, AlphaFold3 ipSAE score, detection in each replicate (Rep1, Rep2, Rep3), number of replicates detected, and subcellular localization. Proteins detected in all three replicates (22 gene loci) are indicated. N Reps indicates the number of replicates in which the representative UniProt accession was detected. Some proteins were detected in ≥2 replicates under alternative UniProt accessions that were merged during gene locus consolidation; see Rep1–3 UniProt ID columns for details.

**Supplementary Table S2. Complete functional enrichment analysis results.** Full results of functional enrichment analysis performed using Metascape (Zhou et al. 2019). Input: 159 genes from the 233 candidate ATM1-proximal proteins that could be mapped to annotated GO terms; background: all *Arabidopsis thaliana* genes. For each enriched term, the GO/KEGG ID, category, description, gene count, percentage, P value, and representative gene symbols are shown.

---

**Supplementary Figure S1.**
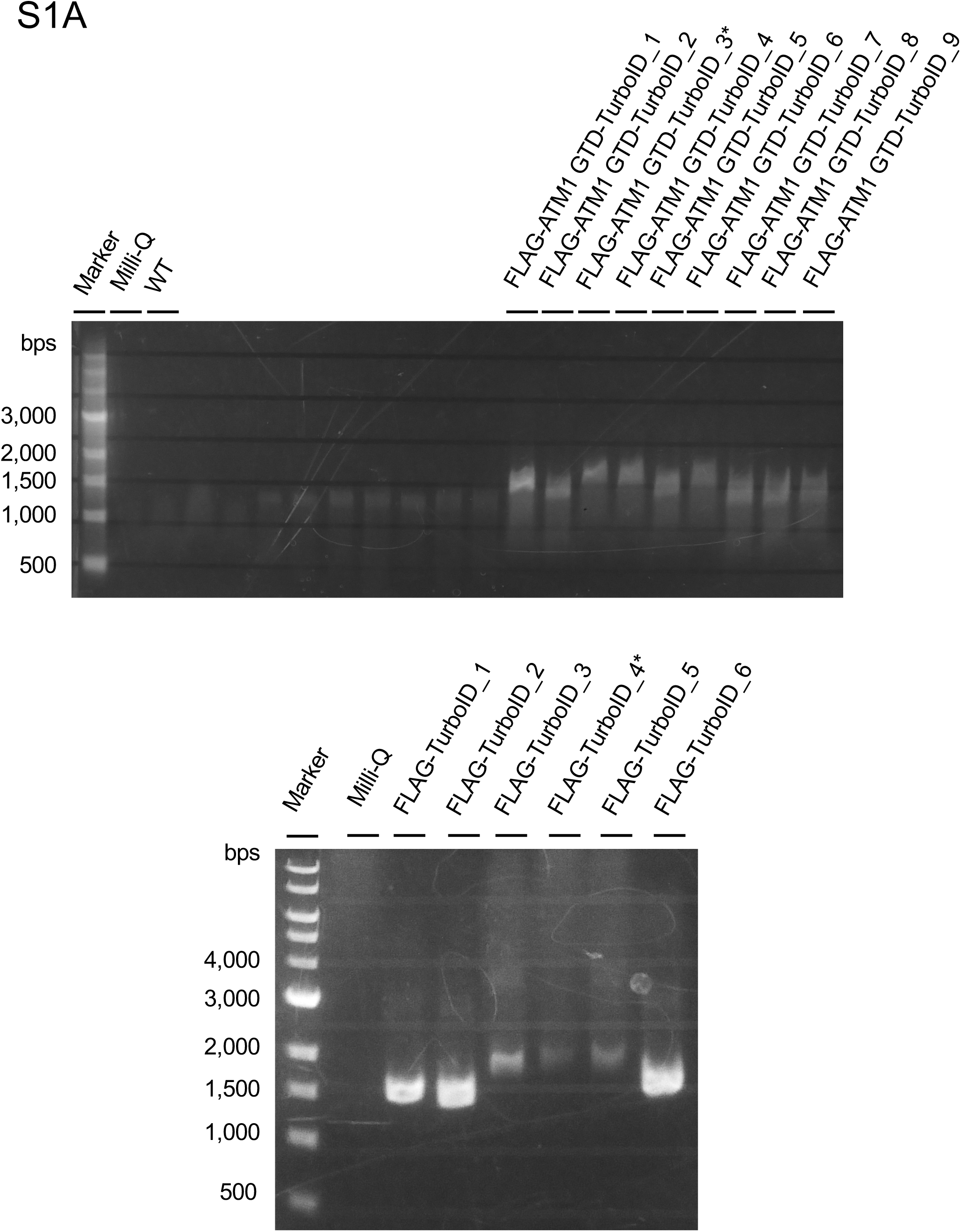
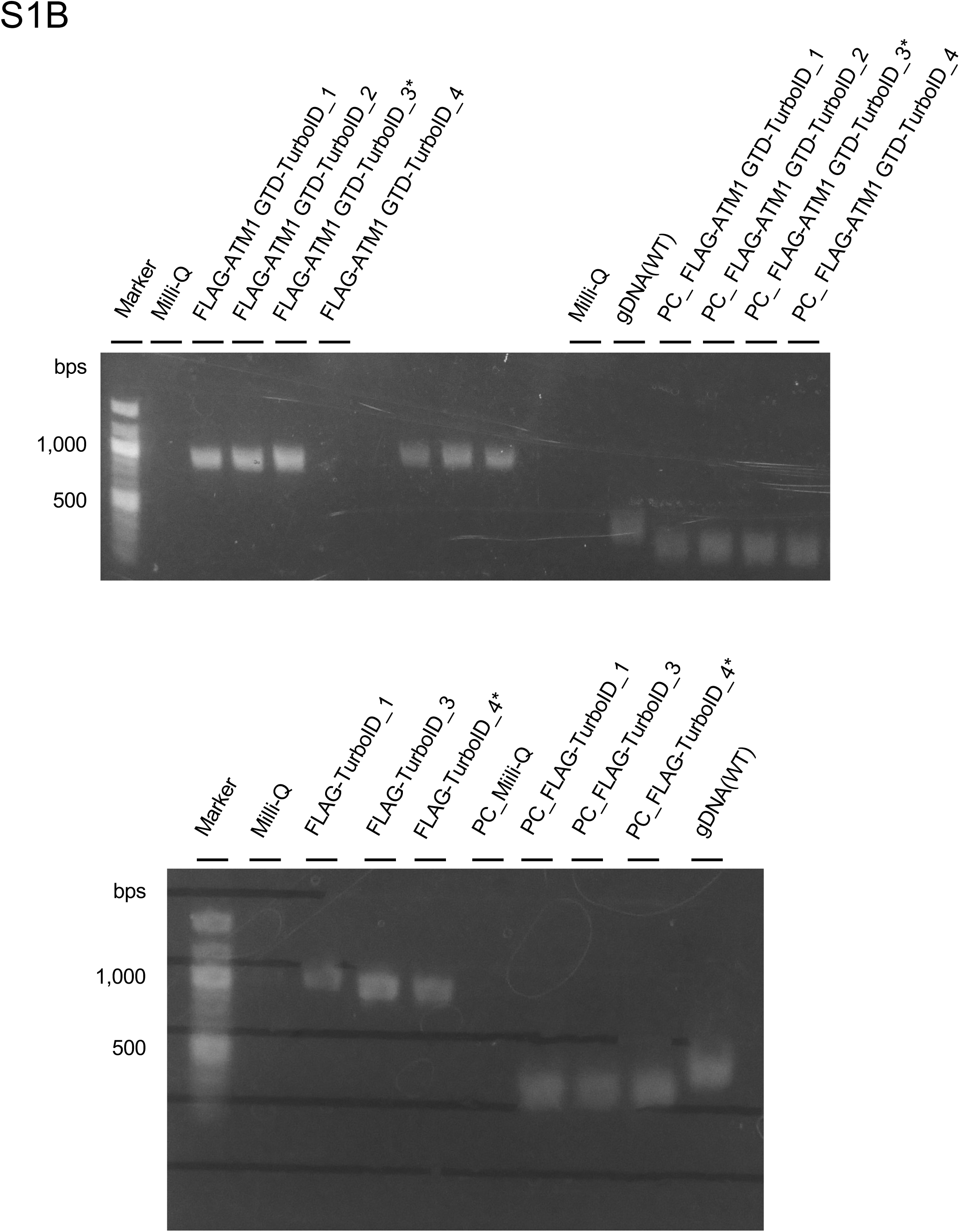
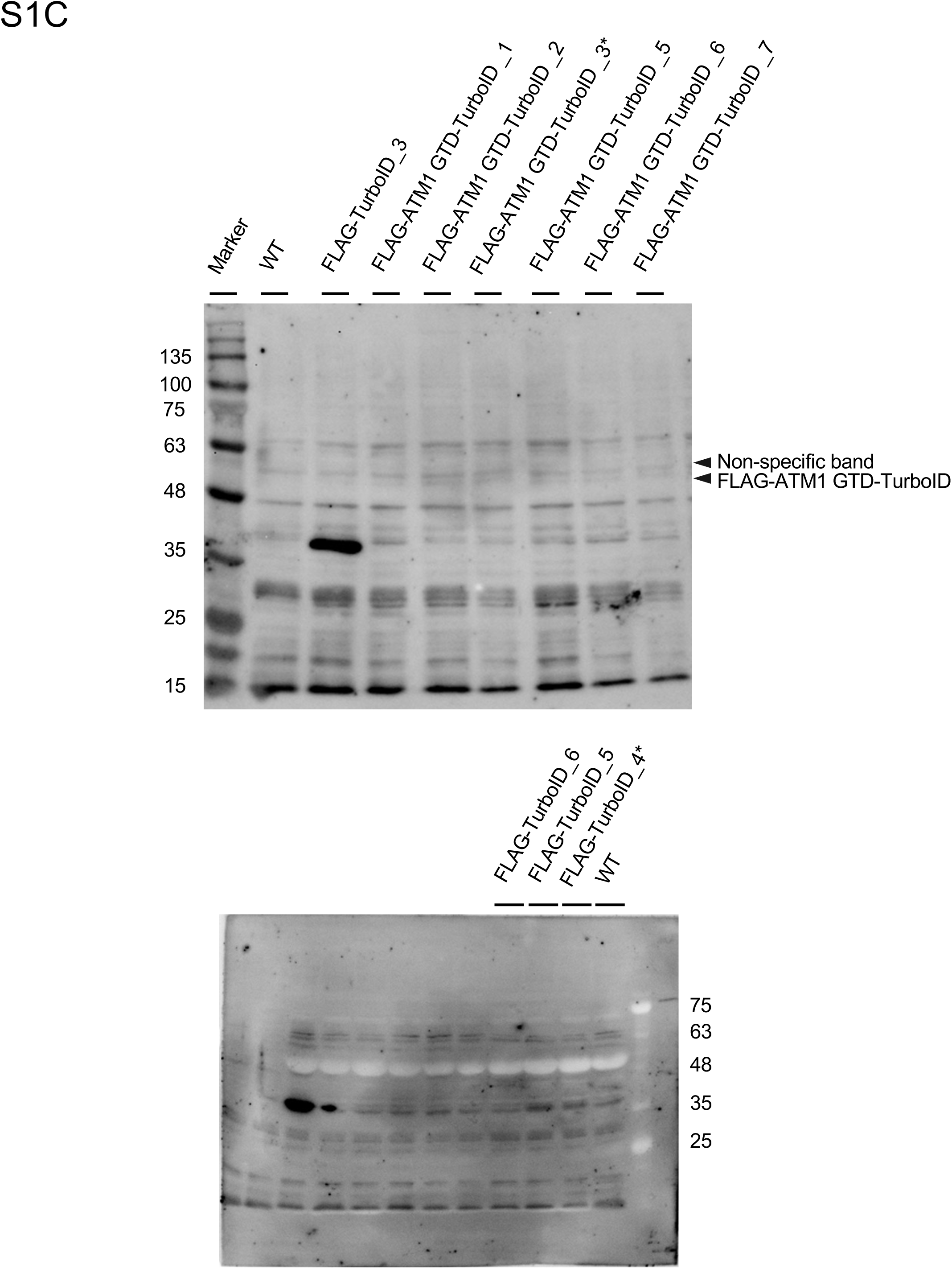
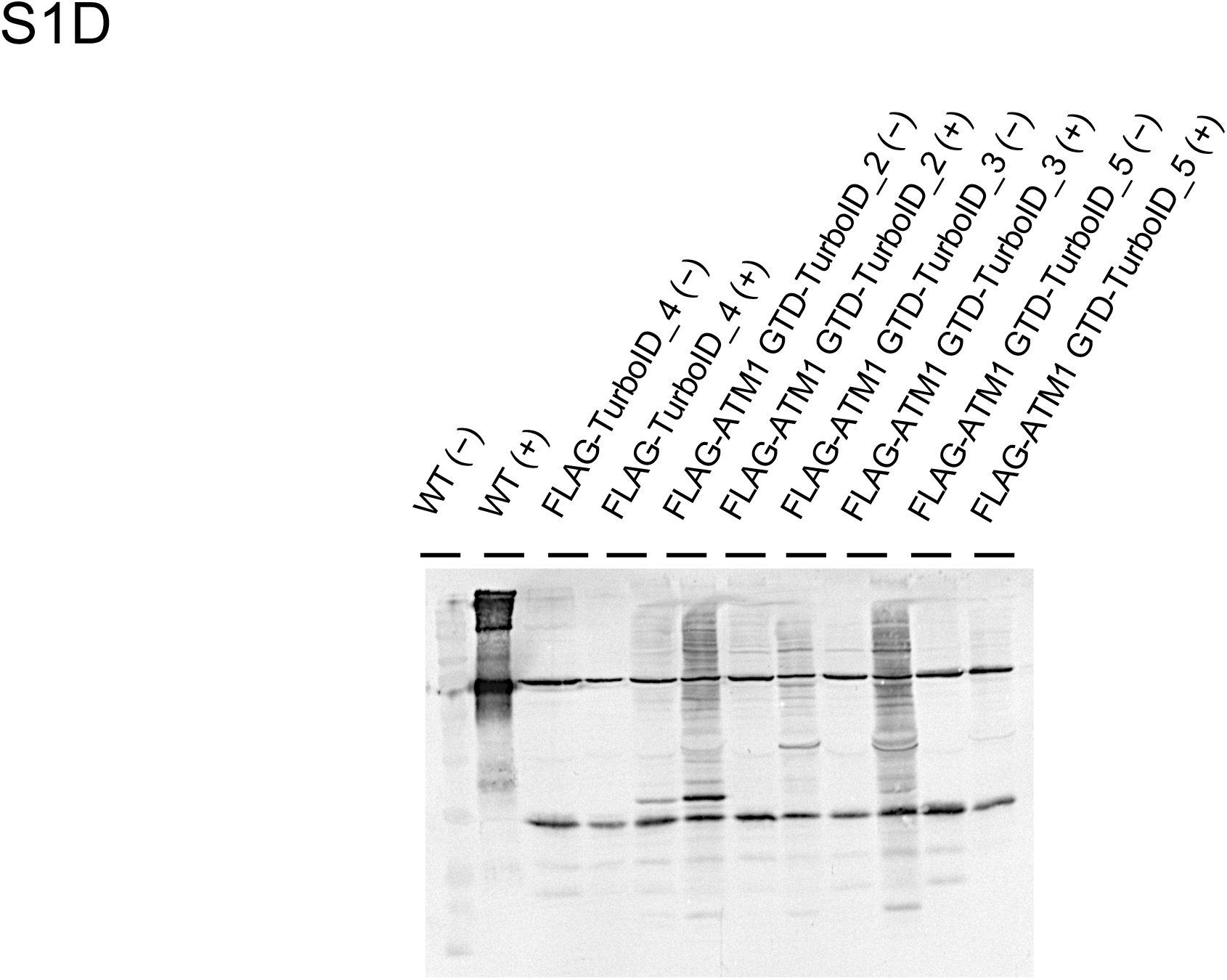
**Characterization of transgenic Arabidopsis plants expressing FLAG-ATM1 GTD-TurboID and FLAG-TurboID.**(A) Genomic PCR analysis confirming transgene integration. Genomic DNA was extracted from independent transgenic lines expressing FLAG-ATM1 GTD-TurboID or FLAG-TurboID (control) and amplified using transgene-specific primers (1,452 bp for FLAG-ATM1 GTD-TurboID; 1,574 bp for FLAG-TurboID). Wild-type (WT) plants and Milli-Q water were used as negative controls. Asterisks indicate lines selected for BioID experiments. (B) RT-PCR analysis confirming transgene expression. Total RNA was extracted from fourteen-day-old seedlings of the indicated lines and reverse-transcribed. Transgene-specific primers were used to detect FLAG-ATM1 GTD-TurboID expression (961 bp). ACTIN2 (ACT2) was amplified as a positive control for cDNA quality (cDNA: 353 bp; gDNA: 439 bp). Genomic DNA (WT) was included as a size control. Asterisks indicate lines selected for BioID experiments. (C) Immunoblot analysis confirming transgene expression. Total protein extracts from fourteen-day-old transgenic seedlings were analyzed by immunoblotting with anti-FLAG antibody (1:2,000). FLAG-ATM1 GTD-TurboID (∼53 kDa) and FLAG-TurboID (∼38 kDa) were detected in the respective transgenic lines but not in wild-type (WT). An open arrowhead indicates a non-specific band. Asterisks indicate lines selected for BioID experiments. (D) Streptavidin-AP blot analysis confirming *in vivo* biotinylation activity. Total protein extracts from fourteen-day-old seedlings treated with (+) or without (−) 500 µM biotin for 3 hours were probed with streptavidin-AP (V559C; Promega). Both FLAG-ATM1 GTD-TurboID and FLAG-TurboID lines confirmed biotinylation activity, demonstrating that TurboID was functional in planta. Asterisks indicate lines selected for BioID experiments.

**Supplementary Figure S2.**
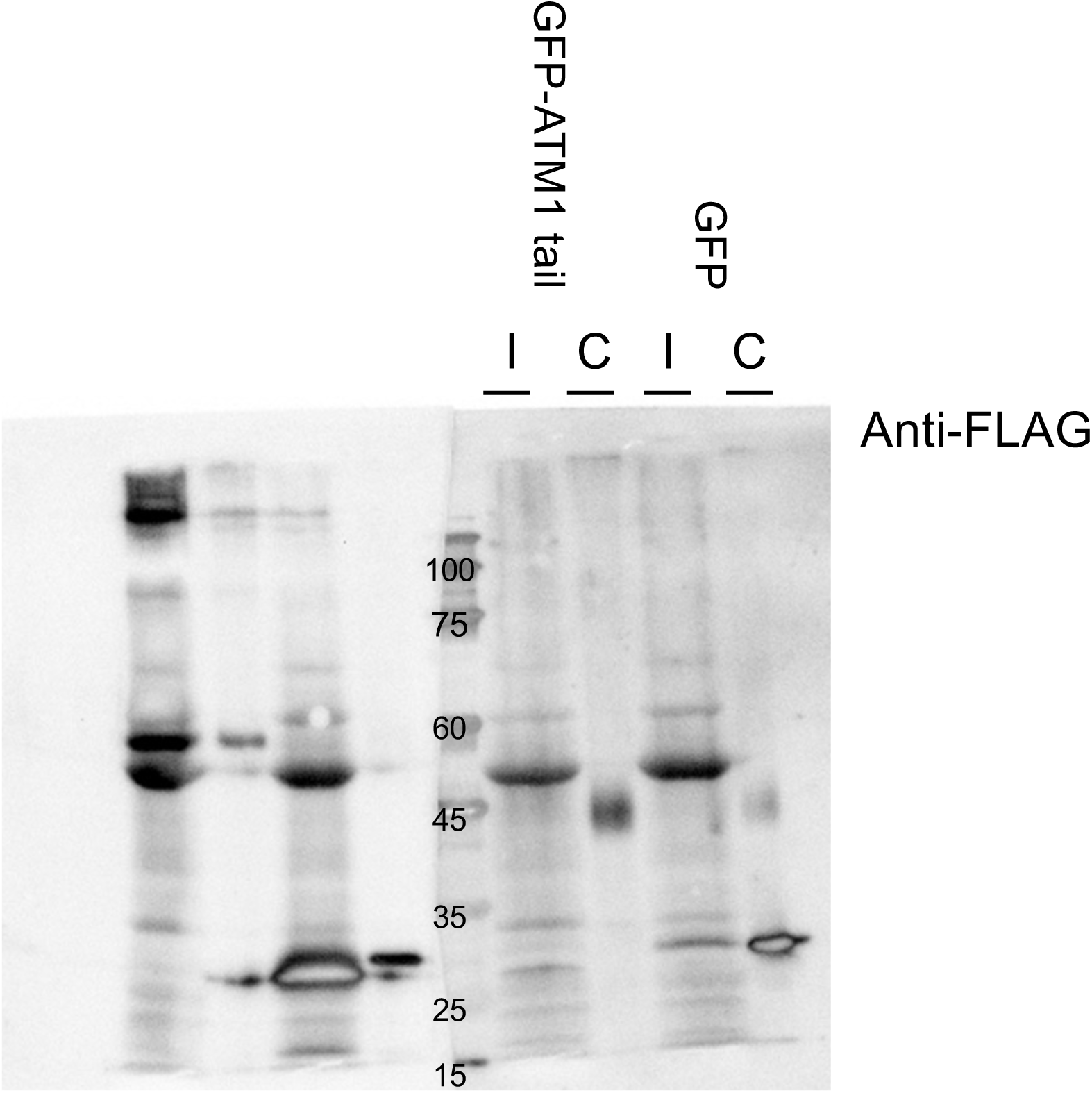
Immunoprecipitation with anti-FLAG beads confirms the identity of the co-precipitated C3H61 band. To confirm that the broad band co-precipitated in Fig. 4A corresponds to FLAG-C3H61, immunoprecipitation was performed using anti-FLAG magnetic beads. GFP-ATM1 tail or GFP alone was co-expressed with FLAG-C3H61 in *N. benthamiana* leaves. Total protein extracts (I, input) and anti-FLAG immunoprecipitated fractions (C, co-IP) were analyzed by immunoblotting with anti-FLAG antibody. FLAG-C3H61 (∼44 kDa) was detected in the co-IP fraction of the GFP-ATM1 tail sample. A faint non-specific band at ∼30 kDa was observed in the GFP control co-IP fraction, which is distinct in size from FLAG-C3H61, supporting the specificity of the co-precipitated band observed in Fig. 4A.

**Supplementary Figure S3.**
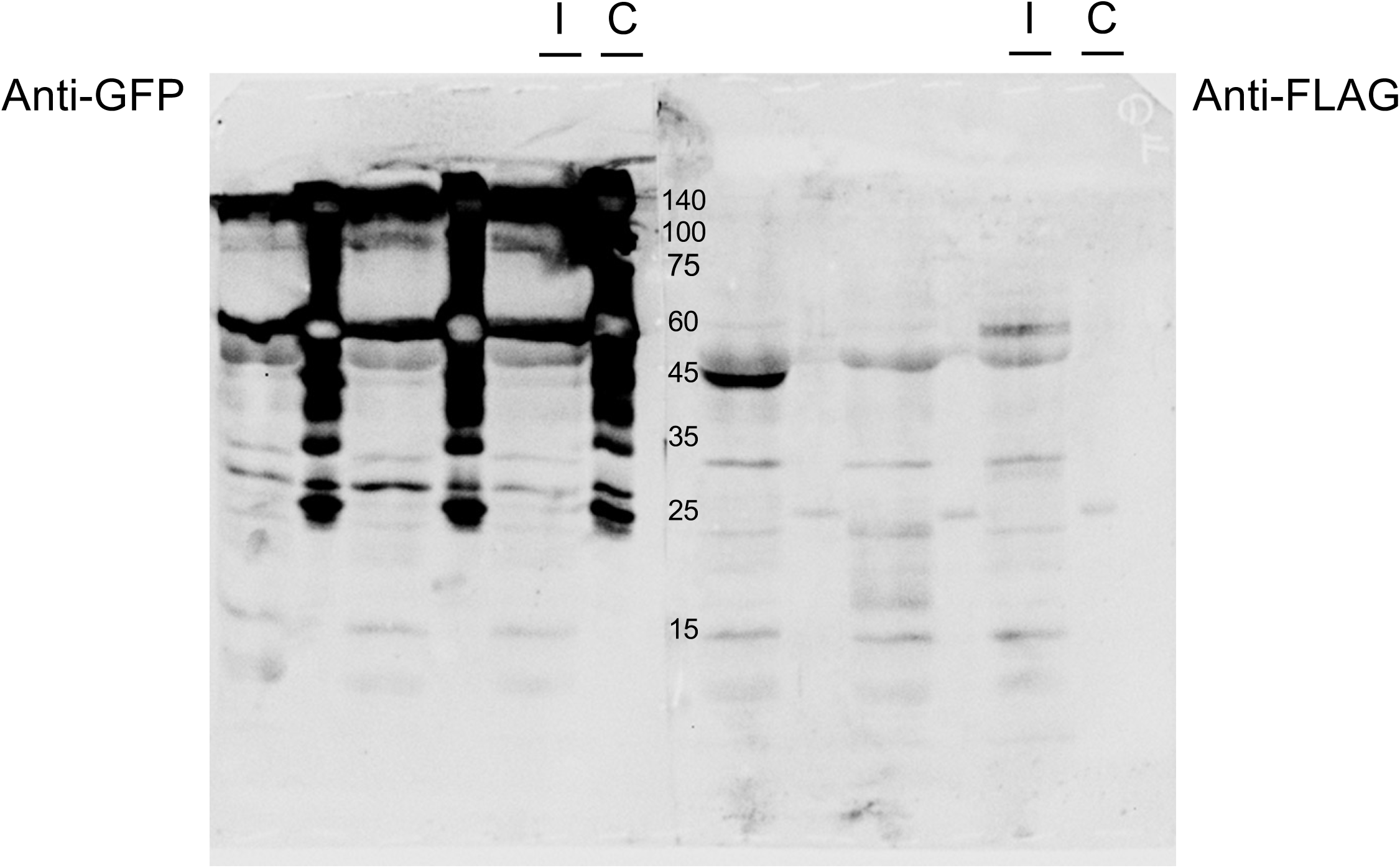
DRB2 shows insufficient expression for co-IP validation. Co-IP assay for DRB2. GFP-ATM1 tail or GFP alone was co-expressed with FLAG-DRB2 in *N. benthamiana* leaves and immunoprecipitated with anti-GFP magnetic beads. Total protein extracts (I, input) and immunoprecipitated fractions (C, co-IP) were analyzed by immunoblotting with anti-GFP (left panel) or anti-FLAG (right panel) antibodies. FLAG-DRB2 expression was insufficient for reliable assessment of the interaction.

**Supplementary Figure S4.**
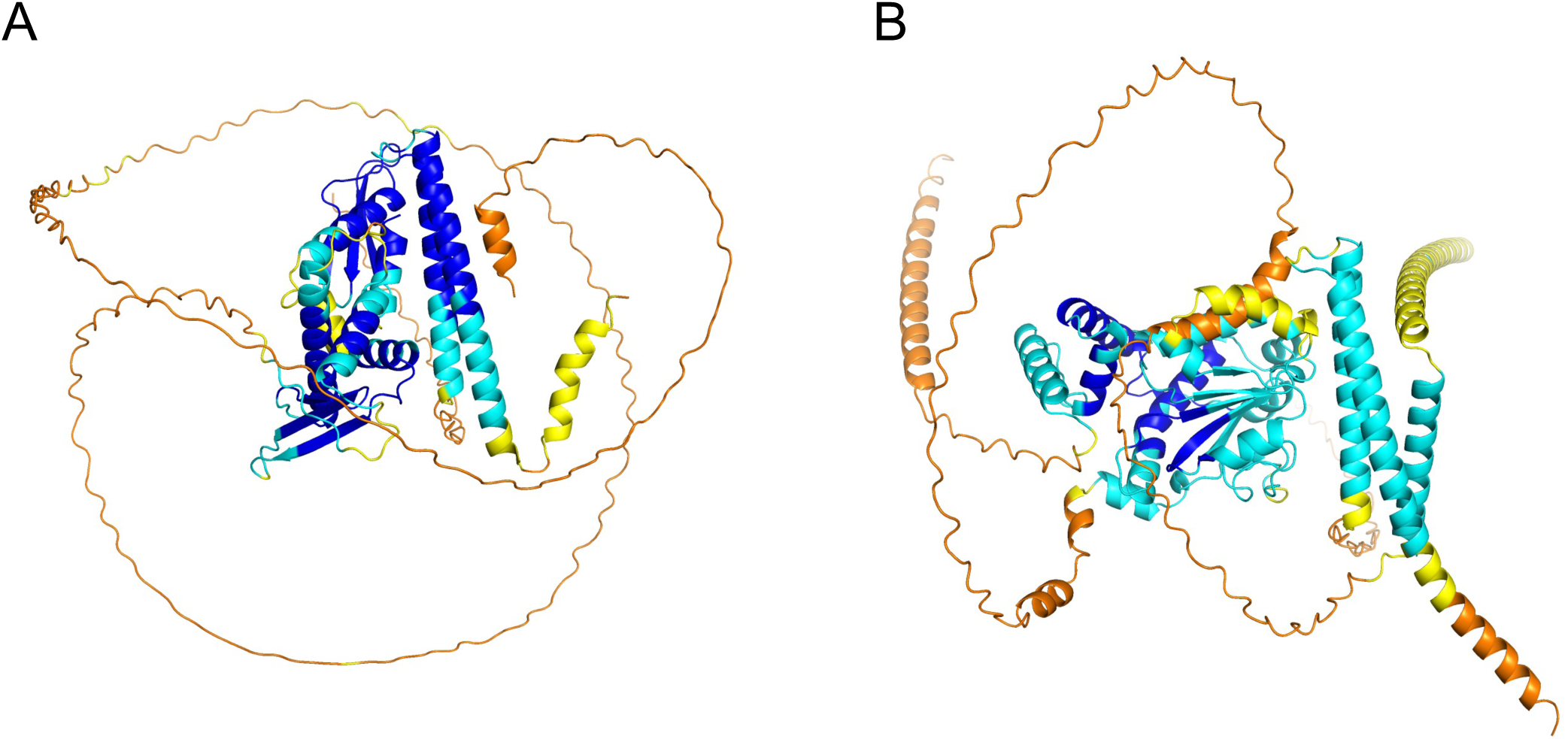
Predicted confidence metrics of AlphaFold3 predicted structures. (A) The prediction confidence (pLDDT) of C-α atoms mapped onto the predicted ATM1–DRB2 complex structure. Blue: 0.9 < pLDDT, Cyan: 0.7 < pLDDT ≤ 0.9, Yellow: 0.5 < pLDDT ≤ 0.7, and Orange: pLDDT ≤ 0.5. The C-termini sequence, located on the right-hand side of the ATM1 GTD’s two alpha helices, was estimated to have low (pLDDT ≤ 0.5) confidence. (B) The confidence score map of the ATM1–SFH7 complex structure. The low-confidence (pLDDT ≤ 0.5) kinked helix structure of SFH7, which is neither Sec14-like nor Nodulin domain, is located in proximity to ATM1.

